# Exploring novel probiotic mechanisms of *Streptococcus* A12 with functional genomics

**DOI:** 10.1101/671420

**Authors:** K Lee, AR Walker, B Chakraborty, JR Kaspar, MM Nascimento, RA Burne

**Affiliations:** Department of Oral Biology, College of Dentistry, University of Florida, Gainesville, Florida, USA; Division of Operative Dentistry, Department of Restorative Dental Sciences, College of Dentistry, University of Florida, Gainesville, Florida, USA

**Keywords:** biofilm ecology, interspecies competition, dental caries, antimicrobial peptides, oral microbiome

## Abstract

Health-associated biofilms in the oral cavity are composed of a diverse group of microbial species that can foster an environment that is less favorable for the outgrowth of dental caries pathogens, like *Streptococcus mutans.* A novel oral bacterium, designated *Streptococcus* A12, was previously isolated from supragingival dental plaque of a caries-free individual, and was shown to interfere potently with the growth and virulence properties of *S. mutans*. Here, we apply functional genomics to begin to identify molecular mechanisms used by A12 to antagonize, and to resist the antagonistic factors of, *S. mutans.* Using bioinformatics, genes that could encode factors that enhance the ability of A12 to compete with *S. mutans* were identified. Selected genes, designated as <u>p</u>otential <u>c</u>ompetitive factors (*pcf)*, were deleted. Certain mutant derivatives showed a reduced capacity to compete with *S. mutans* compared to the parental strain. The A12 *pcfO* mutant lost the ability to inhibit *com*<u>X</u>-inducing <u>p</u>eptide (XIP) signaling by *S. mutans*, while mutants in the *pcfFEG* locus were impaired in sensing of, and were more sensitive to, the lantibiotic nisin. Loss of PcfV, annotated as a colicin V biosynthetic protein, resulted in diminished antagonism of *S. mutans.* Collectively, the data provide new insights into the complexities and variety of factors that affect biofilm ecology and virulence. Continued exploration of the genomic and physiologic factors that distinguish commensals from truly beneficial members of the oral microbiota will lead to a better understanding of the microbiome and new approaches to promote oral health.

**Importance:** Advances in defining the composition of health-associated biofilms have highlighted the important role for beneficial species in maintaining health. Comparatively little, however, has been done to address the genomic and physiological basis underlying the probiotic mechanisms of beneficial commensals. In this study, we explored the ability of a novel oral bacterial isolate, *Streptococcus* A12, to compete with the dental pathogen *Streptococcus mutans*, using various gene products with diverse functions. A12 displayed enhanced competitiveness by: i) disrupting intercellular communication pathways of *S. mutans*, ii) sensing and resisting antimicrobial peptides, and iii) producing factors involved in the production of a putative antimicrobial compound. Research on the probiotic mechanisms employed by *Streptococcus* A12 is providing essential insights into how beneficial bacteria may help maintain oral health, which will aid in the development of biomarkers and therapeutics that can improve the practice of clinical dentistry.

## Introduction

Dental caries is a complex, multifactorial disease that remains the most prevalent chronic disease in children and adults, posing an enormous economic burden to patients and the healthcare system (1, 2). Development of dental caries is a dynamic process in which demineralization of the tooth is driven by repeated acid challenges resulting from the fermentation of dietary carbohydrates by microbes in oral biofilms. Lower pH favors the outgrowth of acidogenic, acid-tolerant species, increasing the proportions of those particular taxa in the microbiota and shifting the microbial ecological balance in the oral cavity (3, 4). With technological advances in high-throughput DNA sequencing, major strides have been made to define the diversity in oral microbiomes associated with dental caries and dental health. A group of caries-associated and health-associated taxa have been identified using metagenomic approaches and a consensus is beginning to develop on which organisms may have the strongest influence on disease development (5–7). Dental caries pathogens belonging to the mutans streptococci group, particularly *Streptococcus mutans*, along with various *Lactobacillus* species, have long been recognized as major contributors to the initiation and progression of caries (8, 9). With the application of metagenomic approaches, the presence of other bacterial taxa, including certain *Bifidobacterium* spp., *Actinomyces* spp., and *Scardovia* spp., as well as *Candida albicans* and certain other fungi (10, 11), have been found to be present in elevated proportions in carious tissues. On the other hand, caries-associated organisms are either absent or their proportions are much lower in health than in disease. Instead, increased proportions of a different group of organisms, including *Streptococcus sanguinis*, *Streptococcus mitis* and *Streptococcus gordonii*, are present in samples from healthy subjects, and the proportions of health-associated taxa decline as the severity of caries increases (12, 13).

Polymicrobial interactions in biofilms are critical determinants of health and disease (14). While *S. mutans* is thought to synergize with other aciduric species to accelerate caries progression, certain microbial interactions interfere with caries pathogens and have the potential to alter the dynamics of oral biofilm behaviors in a way that is beneficial to the host. The overwhelming majority of research on caries microbiology has been focused on understanding the pathogenicity of species driving caries progression. However, a body of evidence is accumulating that supports that oral bacteria that are associated with health actively promote oral health by employing various strategies that work directly or indirectly to foster an environment that is less favorable for the establishment and persistence of caries pathogens, including *S. mutans*. Some antagonistic strategies employed by commensal streptococci and beneficial bacteria are well defined. For example, H_2_O_2_ production by commensal oral streptococci appears to have a substantial impact on oral biofilm ecology (15); H_2_O_2_ inhibits the growth of *S. mutans* and many other oral pathogens at concentrations that do not appreciably affect the producing strains. In addition, the enzymatic alkalinization of oral biofilms by oral bacteria, especially through the breakdown of arginine by the arginine deiminase system (ADS), is a critical contributor to pH homeostasis in oral biofilms. Alkali production can prevent the outgrowth of caries-causing pathogens and can shift the chemical balance in favor of tooth remineralization (16, 17). Similarly, urea is secreted in millimolar quantities in saliva and can be metabolized by bacterial ureases in oral biofilms to yield ammonia, elevating biofilm pH. Both the ADS and urease release from their respective substrates one molecule of CO_2_ and two of ammonia, which can protect acid-sensitive bacteria from growth inhibition or killing at low pH (18, 19) by raising the cytoplasmic pH, which has bioenergetic benefits, and increasing the pH of the environment. Utilization of arginine by the ADS is particularly advantageous because it also generates ATP, which can be used by ADS-positive organisms for growth and maintenance (20). A group of oral streptococci that commonly comprise a substantial proportion of the oral microbiome (21) express the ADS, including the highly arginolytic clinical isolate *Streptococcus* A12 (22). *Streptococcus* A12 was isolated from supragingival plaque of a caries-free subject and has the ability to moderate biofilm pH and potently interfere with the growth and virulence-related properties of *S. mutans*. Compared to the highly arginolytic reference strain *S. gordonii* DL1, A12 was shown to have higher arginine deiminase (AD) enzyme activity and much less arginine was required to induce significant levels of AD activity. A12 inhibits the growth of *S. mutans* through robust pyruvate oxidase (Pox) -dependent H_2_O_2_ production, expressing substantially higher Pox activity than *S. gordonii* DL1. Importantly, A12 was able to interfere with CSP (competence-stimulating peptide)-mediated activation of bacteriocins (mutacins) by *S. mutans* through a protease encoded by the *sgc* called Challisin, similar to *S. gordonii* DL1 (23). However, unlike *S. gordonii*, A12 also efficiently inhibited XIP-dependent signaling through an undefined pathway (22).

The oral cavity presents tremendous opportunities for unraveling the complexities of microbial communities in humans, but also an immense challenge as it is one of the more heterogeneous microbial ecosystems in the human body. Metagenomic analyses of oral microbial communities has facilitated the identification of the microbes that may most profoundly influence health and disease (7, 21, 24), although inter-subject variability in microbiome composition renders generalizing about the contribution of specific taxa challenging. Understanding the etiology of oral diseases is further complicated by the relatively recent demonstration that individual isolates of many abundant oral bacterial species display tremendous genomic and phenotypic heterogeneity. Initially, it was established that there was a high degree of genomic diversity in the dental caries pathogen *S. mutans* (25), and that the genomic diversity was correlated with similarly substantial diversity in a variety of phenotypic properties that are related to the pathogenic potential of the organisms (26). More recently, it was established that high levels of genomic diversity and variability in selected beneficial properties exist among the oral commensal streptococci that are the most abundant members of the oral microbiome (27, 28). Clearly, then, sequencing of DNA or RNA from human samples cannot alone predict how members of the oral microbiome function and interact with one another, and with their host, to determine the pathogenic potential of oral biofilms. Functional genomics combined with various model systems will need to be an integral part of the effort to develop a more comprehensive understanding of the spectrum of mechanisms used by beneficial bacteria to exert probiotic effects, with the ultimate goal being to ameliorate oral health by modulating the composition and activity of the oral microbiome. Here, we employ functional genomics to begin to dissect mechanisms by which the beneficial organism *Streptococcus* A12 may suppress the growth and virulence of the dental caries pathogen *S. mutans*.

## Materials and Methods

### Bacterial strains and growth conditions

Table 1 lists all strains and plasmids used in this study. *Streptococcus* A12 was isolated from supragingival plaque of a caries-free subject (22). A12, *S. mutans* UA159 and genetically modified derivatives of each strain were routinely grown in brain heart infusion (BHI) broth, on BHI agar plates (Difco), or in the chemically defined medium FMC (29) at 37°C in a 5% CO_2_ aerobic atmosphere. To select for antibiotic-resistant A12 strains after genetic transformation or to enumerate *S. mutans* in dual-species cultures, BHI plates containing kanamycin (1 mg ml^−1^) were used.

**Table 1.** Bacterial strains used in this study

| Table 1 Bacterial strains used in this study |  |  |
| --- | --- | --- |
| Strains | Relevant genotype | Source |
| A12 | Isolated from supragingival dental plaque of caries-free subject | Clinical strain, 22 |
| A12ΔpcfV | A12ΔpcfV; Km<sup>r</sup> | This study |
| A12ΔpcfO | A12ΔpcfO; Km<sup>r</sup> | This study |
| A12ΔpcfFEG | A12ΔpcfFEG; Km<sup>r</sup> | This study |
| A12ΔpcfRK | A12ΔpcfRK; Km<sup>r</sup> | This study |
| A12Δ1179 | A12Δ1179; Km<sup>r</sup> | This study |
| A12Δsgc | A12Δsgc; Erm<sup>r</sup> | 22 |
| A12ΔspxB | A12ΔspxB; Km<sup>r</sup> | 22 |
| *S. gordonii* DL1 | Wild-type reference strain | Laboratory stock |
| *S. sanguinis* SK150 | Wild-type reference strain | Laboratory stock |
| *S. mutans* UA159 | Wild-type reference strain | ATCC |
| *S. mutans* pBGS | *S. mutans* UA159:: pBGS; Sp<sup>r</sup> | This study |
| SAB358 | *S. mutans* UA159::PcomX-lacZ; Km<sup>r</sup> | 35 |
| SAB249 | *S. mutans* UA159::PcipB-lacZ; Km<sup>r</sup> | 22 |

| Plasmids |  |  |
| --- | --- | --- |
| pBGS | *Streptococcus* integration vector, Sp<sup>r</sup> | 34 |

Km<sup>r</sup>, kanamycin resistance; Sp<sup>r</sup>, spectinomycin resistance; Em<sup>r</sup>, erythromycin resistance

*S. mutans* synthetic CSP (sCSP), corresponding to the 21-aa peptide (30) was synthesized by the Interdisciplinary Center for Biotechnology Research (ICBR) facility at the University of Florida, and its purity (97%) was confirmed by high-performance liquid chromatography (HPLC). Synthetic XIP (sXIP), corresponding to residues 11 to 17 of ComS {aa sequence GLDWWSL, (31)} was synthesized and purified to 96% homogeneity by NeoBioSci (Cambridge, MA). To synthesize A12 CSP to be utilized to induce competence in A12, the CSP precursor amino acid sequence (ComC) located upstream of ComDE was identified and an active 17-amino acid peptide CSP was predicted to be derived from ComC following the cleavage near a conserved double glycine (GG) cleavage motif. The 17-aa A12 CSP was synthesized by Biomatik (96.8% purity reported by the supplier).

### Construction of mutant strains and reporter gene fusions

Mutant strains of A12 were engineered by double-crossover recombination using linear DNA assembled through a Gibson Assembly^®^ kit (New England Biolabs, Beverly, MA) (32). Briefly, primer sets (Table 2) were designed to PCR amplify two DNA fragments flanking the coding sequence of the genes of interest and containing at least 25 bases of sequence that overlapped with the termini of the non-polar kanamycin-resistance cassette in pALH124 (33). The two flanking DNA fragments and the kanamycin cassette were mixed in equimolar concentrations in a single isothermal ligation reaction. Overnight cultures of A12 were inoculated into fresh BHI cultures and the ligated DNA products (0.5 µg) were used to transform A12 in BHI using 50 nM A12 sCSP to induce competence. After 3 h of incubation, cells were plated onto BHI agar with Km (1 mg ml^−1^) and isolated colonies were picked for PCR verification. PCR products from positive transformants were sent for DNA sequencing to ensure that the fragment was inserted into the desired locus and that no mutations were present in the flanking regions used for homologous recombination. To construct a marked strain for dual-species experiments, *S. mutans* was transformed with pBGS, which allowed for stable integration of a spectinomycin marker into the *gtfA* gene (34), yielding the spectinomycin resistant (Sp^r^) *S. mutans* strain. Construction and characteristics of strains of *S. mutans* carrying a *lacZ* gene fused to the promoter of *cipB* (P*_cipB_*::*lacZ)* or *comX* (P*_comX_::lacZ)* are detailed elsewhere (22, 35).

**Table 2.**
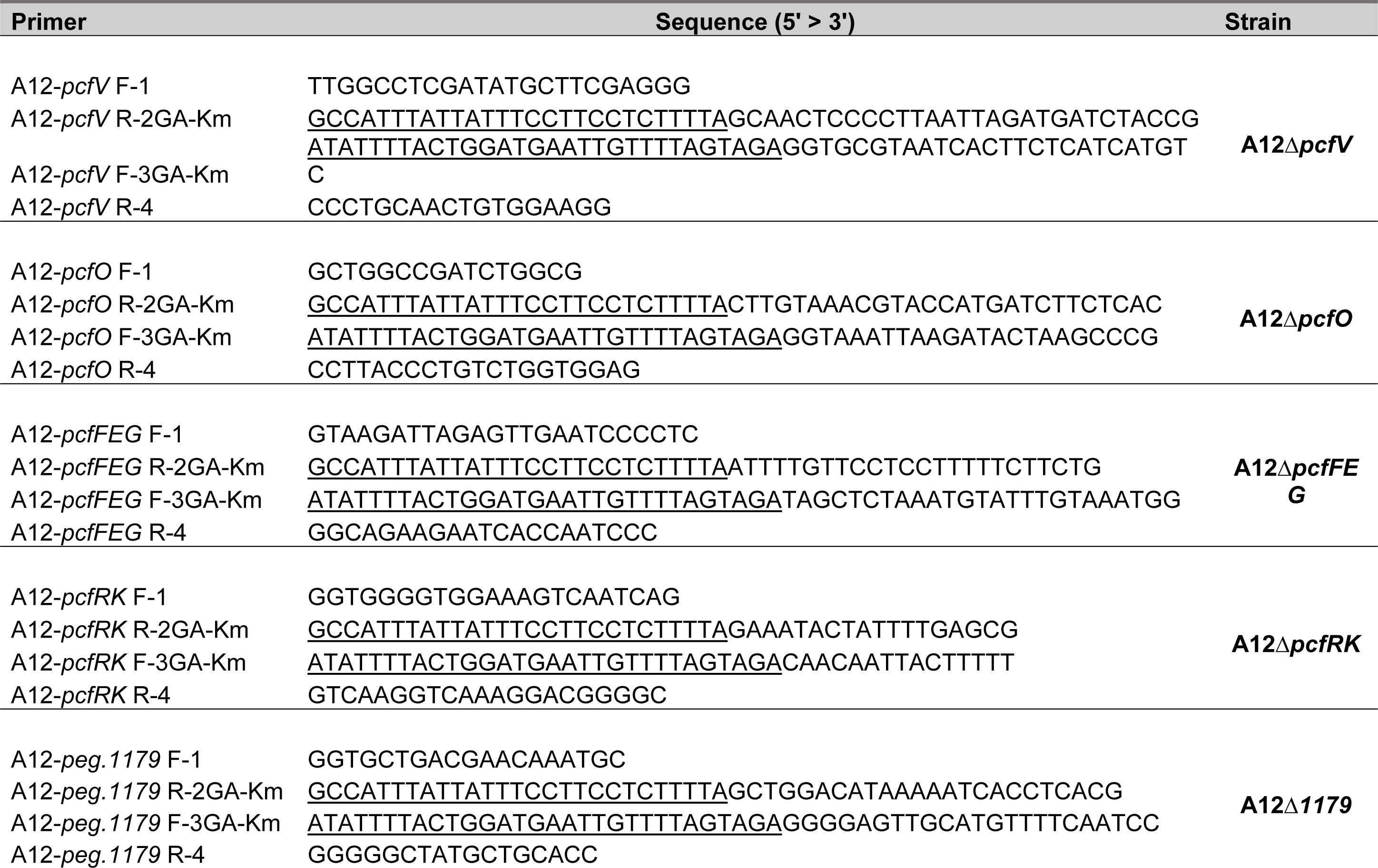
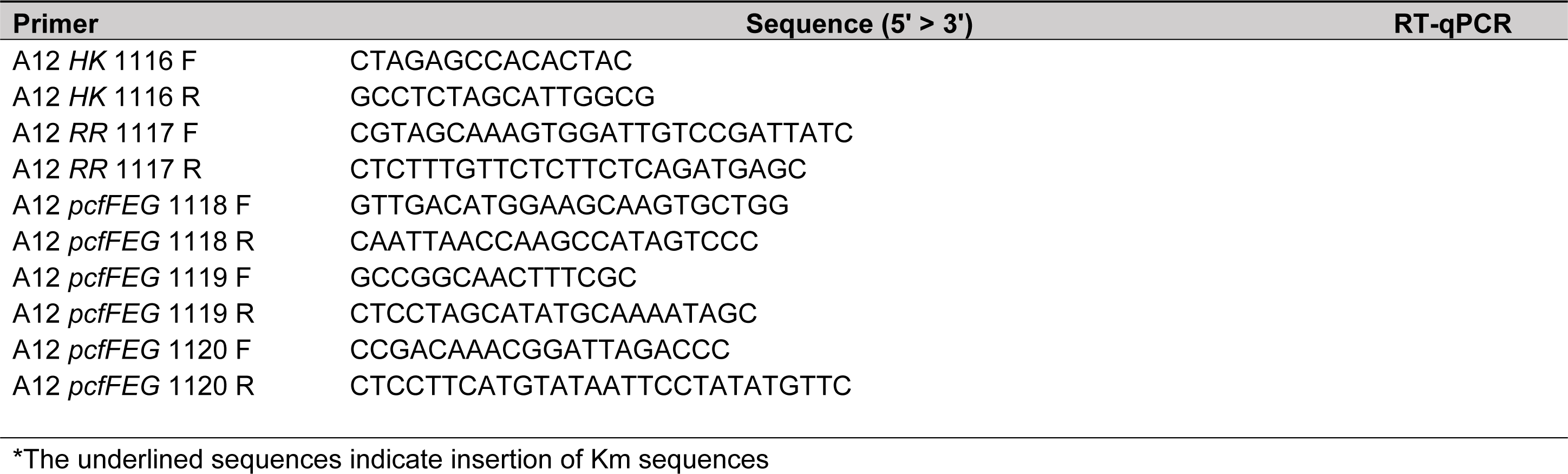
Primers used in this study

### Monitoring of growth

Growth of A12 and A12 mutant derivates was monitored using a Bioscreen C (Growth Curves USA, Piscataway, NJ). Overnight cultures were diluted 1:20 in BHI and were grown to an OD_600_ of 0.5. Subsequently, the culture was diluted 1:100 into BHI medium with or without nisin and transferred to 100-well honeycomb plates. The OD_600_ was measured every 20 minutes, with shaking for 10 s before each measurement. When noted, sterile mineral oil was overlaid in each well to reduce the exposure of cells to oxygen.

### Antagonism assays on agar plates

A plate-based antagonism assay was utilized to detect and measure growth inhibition between *S. mutans* UA159 and A12, or A12 derivatives (22). Briefly, bacterial cultures were grown overnight in BHI medium, then the OD_600_ was adjusted to 0.5. Aliquots (6 µl) from each culture were spotted adjacent to the other strain on BHI agar plates, either simultaneously or with A12 strains inoculated first, followed by inoculation with *S. mutans* 24 h later. All plates were incubated for 24 or 48 h at 37°C in a 5% CO_2_ aerobic atmosphere. Zones of inhibition were captured with a digital imager and quantified with NIH Image J analysis, which was set to standardized scale (10.234 pixels per mm). Deferred antagonism assays were also performed to monitor the effects of A12 and its derivatives on bacteriocin production by *S. mutans.* Cultures of Sp^r^ *S. mutans* (Table 1), *S. sanguinis* SK150 and A12 were grown overnight, centrifuged at 10,000 × *g* and washed with PBS. The cells were resuspended in PBS and adjusted to OD_600_ = 0.5. *S. mutans* was then mixed with other strains in a 1:1 ratio, based on optical density. Cultures of *S. mutans* or A12 alone were used as controls. Single or mixed cultures were stabbed onto BHI agar with a sterile toothpick and grown in aerobic (5% CO_2_) or anaerobic (GasPak^TM^, BD Life Sciences) conditions at 37°C. After 24 h of incubation, 3 ml of soft agar overlay (BHI in 1% agar) containing 10^7^ cells of the indicator strain *S. sanguinis* SK150 was poured evenly onto the plate. All plates were incubated for an additional 24 h and zones of inhibition were measured from digital images. To enumerate CFUs after 48 h incubation, agar plugs were obtained from the center of the zone, resuspended in 1 ml of PBS, and sonicated for three 30-second cycles in a sonicating water bath, with cooling on ice during intervals. Samples were then serially diluted and plated onto BHI agar or BHI agar containing spectinomycin (1 mg ml^−1^) or kanamycin (1 mg ml^−1^).

### Monitoring of *comX* or *cipB* gene promoter activity

*S. mutans* UA159 carrying a *lacZ* gene from *Streptococcus salivarius* fused to the promoter regions of *comX* (encodes for the alternative sigma factor required for genetic competence) or *cipB* (encodes for a bacteriocin) (36) were utilized to assess the expression of those genes in the presence of XIP or CSP, as described elsewhere (22, 35). To determine whether *S. gordonii* DL1 or strains of A12 had the ability to interfere with XIP or CSP signaling of *S. mutans*, supernates from cultures grown in FMC medium (for XIP signaling) or BHI medium (for CSP signaling) were collected by centrifugation, the pH of the supernatant fluid was adjusted to 7.0, and, in the case of FMC medium for XIP signaling, additional glucose was added to increase the concentration by 25 mM. All supernates were then filtered through a 0.2 µm syringe filter. Similarly prepared *S. mutans* UA159 supernates were included as a positive control. Various concentrations of sXIP were then added to supernates and incubated overnight at 37°C. Cultures of the reporter strain *S. mutans* UA159::P*_comX_-lacZ* grown in FMC medium to an OD_600_ of 0.15 were then incubated with the supernates for 2 h and cells were collected to assay β-galactosidase (LacZ) activity as an indicator of *comX* promoter activity. For CSP signaling, 2 µM CSP was added to the supernates and incubated for 3 hours. Reporter strain *S. mutans* UA159::P*_cipB_-lacZ* grown in BHI medium to an OD_600_ of 0.15 was then incubated with the samples for an additional 2 hours. Cells were then collected for LacZ activity as a measure of *cipB* promoter activity. β-galactosidase activity was determined using a modification of the Miller protocol (22, 37). Briefly, cells were harvested by centrifugation, washed once with Z buffer (Na-phosphate buffer pH 7.0, 10 mM KCl, 1 mM MgSO_4_, 5 mM β-mercaptoethanol) and resuspended in Z buffer. A sample aliquot was treated with toluene-acetone (1:9) for 2 minutes, and the remainder of each sample was used to measure OD_600_. The reaction was initiated by adding 80 µl of a 4 mg/ml solution of ONPG (*o*-nitrophenyl-β-D-galactopyranoside) and terminated by adding 500 µl of 1M Na_2_CO_3_. Samples were then quickly centrifuged at 15,250 × *g* for 1 minute and the optical density of the supernatant fluid was measured at 420 and 550 nm. All assays were performed using biological triplicates, with technical triplicates for each sample. β-galactosidase activity is reported in Miller units.

### Analysis of gene expression by quantitative reverse-transcriptase PCR (qRT-PCR)

Overnight cultures of A12 strains were inoculated into 7 ml of fresh BHI medium. Once cultures reached an OD_600_ of 0.5, cultures were treated with or without 2 µg/ml nisin and incubated for 15 min at 37°C. Cells were then harvested by centrifugation (5 min, 10,000 × *g*) and immediately suspended in 1 mL bacterial RNA protect (Qiagen, Germantown, MD), followed by 5 min incubation at room temperature. Total RNA was extracted as previously described (38, 39) and 1 µg of total RNA was reverse transcribed to cDNA using the iScript Select cDNA Synthesis Kit (Bio-Rad) with random hexamers. Total RNA and cDNA were prepared in biological triplicates and subsequent technical triplicates for each cDNA template were included with appropriate controls. The obtained cDNAs were used as templates for quantitative real-time PCR (qRT-PCR) for gene expression analysis. The levels of various mRNA transcripts were quantified using gene-specific primers (Table 2) and normalized to 16S rRNA transcripts. qRT-PCR was performed using an iCycler iQ real-time PCR detection system (CFX96;Bio-Rad) and iQ SYBR^®^ green Supermix (Bio-Rad), according to the protocols provided by the supplier.

### Pangenome analysis on isolates closely related to A12

Bioinformatic comparisons of recently isolated strains that were genetically similar to A12 (28) were conducted in two steps. First, the databases required for the analyses were built. The annotations for all the strains considered in this study were generated with the open source software Prokka (40). The protein databases for Blast searches (41, 42) were built from the Prokka translated coding gene output. Lastly, the pangenome analysis was conducted with the Roary pipeline (43), which clustered the isolates based on gene presence. Phylogenetic maximum likelihood trees from multiple sequence alignments from Roary were built using Phyml (44, 45) and annotated, then displayed with iTol (46).

## Results

### A12 genes that enhance competition with *S. mutans*

Using various bioinformatic tools (e.g. Blast, GenBank, Patric), a discrete set of genes that encode factors that might augment the ability of A12 to compete with *S. mutans* were identified and designated as *pcf* genes, for <u>p</u>otential <u>c</u>ompetitive factors. This study focused on three genetic loci (Fig. 1A). The *pcfO* gene (ATM98_08220) was identified based on its homology to a protease reported to disrupt quorum sensing in *Streptococcus pyogenes* (47) and the gene clusters containing *pcfFEG* (ATM98_05540, ATM98_05535, ATM98_05530) and *pcfV* (ATM98_05815) were chosen due to their association with antimicrobial peptides. The *pcfFEG* genes encode a predicted three-component ABC transporter that consists of two permeases (ATM98_05535 and ATM98_05530) and a protein with the Walker motif ATP binding domain (ATM98_05540). The product of *pcfV* is annotated as a colicin V biosynthetic protein. To determine whether the products of these genes could influence the ability of A12 to compete with *S. mutans*, a collection of mutant strains was generated (Table 1) using double cross-over homologous recombination in which the genes of interest were replaced with a non-polar kanamycin resistance (Km^r^) marker. A12 and the mutants were tested in plate-based growth inhibition assays either with A12 spotted first, followed by *S. mutans* 24 h later, or when A12 and its derivatives were spotted simultaneously with *S. mutans*, as detailed in the methods section. As previously reported, wild-type A12 was able to inhibit the growth of *S. mutans* as evinced by a large zone of inhibition when spotted first (Fig. 1B), and to a lesser extent when spotted simultaneously. When A12 strains were inoculated prior to *S. mutans*, deletion of *pcfFEG, pcfV* and *pcfO* resulted in significantly less antagonism of *S. mutans* compared to wild-type A12, although the effect of deletion of *pcfO* was less profound (Fig.1C). When A12 or its derivatives were spotted simultaneously with *S. mutans* UA159, the ability of all strains to inhibit *S. mutans* was similar. The phenotypes observed were not due to any substantial growth defects as the growth rates of the mutants were generally comparable to the parental strain, as were the maximum optical densities attained and the optical densities attained after 16 h of growth (Fig. S1).

**Figure 1.**
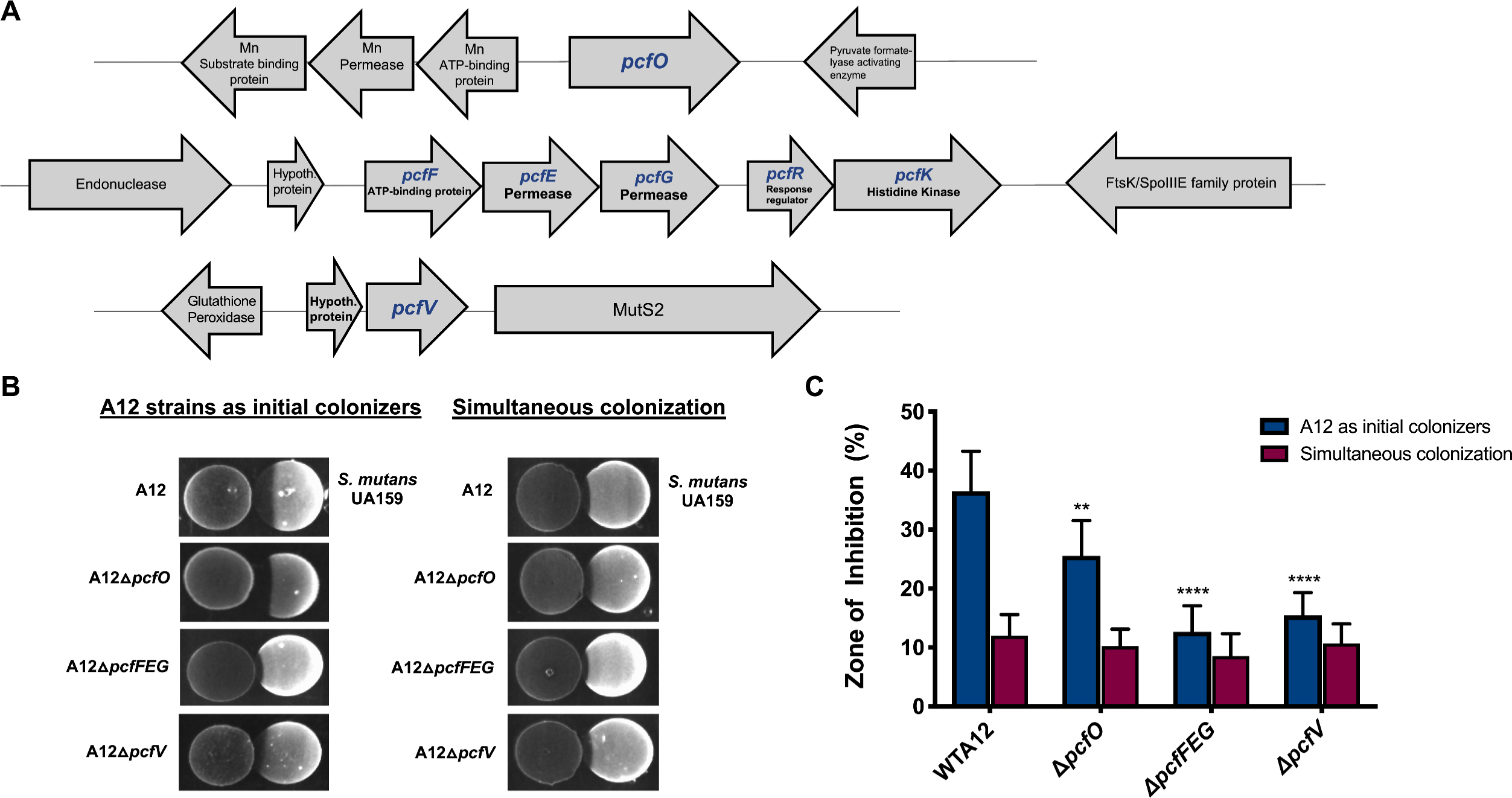
(A) Schematic diagram of loci encoding <u>p</u>otential <u>c</u>ompetitive factors of A12. *pcf* genes are indicated in blue and illustrated are gene order and the arrangement of neighboring genes. (B) Plate-based growth inhibition of A12 strains versus *S. mutans*. Bacterial cultures were grown overnight in BHI medium and adjusted to an OD_600_ of 0.5 with sterile BHI. Aliquots (6 µl) from each culture was spotted adjacent to the other strain on BHI agar plates whether simultaneously (right), or A12 or A12 mutant derivatives first, followed by spotting of *S. mutans* 24 h later (left). Plates were incubated for 24 h or 48 h, respectively, at 37°C in a 5% CO_2_ aerobic atmosphere. Representative images are shown. (C) Zones of inhibition were captured with a digital imager and measured with NIH Image J analysis, which was set to standardized scale (10.234 pixels per mm). The area of inhibition was divided by the total area of the respective bacterial spot in millimeters × 100 to yield % Zone of Inhibition. Values are the average of two biological replicates, performed in technical quadruplicates. Error bars represent standard deviations. Asterisks indicate statistically significant differences compared to the zone of inhibition created by wild-type A12. Statistical analysis was performed using an unpaired Student’s t-test; ** P < 0.01, **** P < .0001.

### A12 mutant strains have an altered ability to tolerate antagonistic factors of *S. mutans*

To further assess the impact of the *pcf* genes in the antagonism of *S. mutans*, interaction between A12 or A12 mutant strains and *S. mutans* was evaluated using a dual-species, agar plated-based system in which A12 and *S. mutans* were first cultured separately in planktonic culture to mid-exponential phase. The cultures were mixed in a 1:1 ratio based on optical density and inoculated onto BHI agar by gently stabbing with a toothpick that had been dipped into the mixed culture. After 24 hours of incubation, the viable proportions of A12 and *S. mutans* were determined by aseptically obtaining the agar plugs and dispersing the bacteria. Controls included pure cultures of A12 or *S. mutans* grown, inoculated into plates, and enumerated in the identical manner. When the plates were incubated in aerobic conditions, almost no *S. mutans* cells were recovered when co-cultivated with A12 or any of the A12 *pcf* mutant derivates (Fig. 2A). Since a primary mechanism for commensals, including A12, to antagonize *S. mutans* in aerobic conditions is through the production of H_2_O_2_ (22, 27), an *spxB* mutant of A12 (22) was included to estimate the relative contribution of H_2_O_2_ to the observed growth inhibition; *spxB* encodes the H_2_O_2_-producing pyruvate oxidase that is the primary source of H_2_O_2_ produced by this organism (22). Compared to co-cultures with wild-type and *pcf* mutants of A12, *S. mutans* constituted 38% of the population of viable cells recovered from the plates when co-inoculated with the *spxB* mutant and incubated under aerobic conditions. Under anaerobic conditions, A12 strains lacking *pcfO*, *pcfFEG*, or *pcfV* were less competitive with *S. mutans* compared to wild-type A12, with significantly lower proportions of CFUs of A12Δ*pcfO* and A12Δ*pcfFEG* recovered after 24 hours. Also of note, there were no differences in the CFUs of pure cultures of A12 in anaerobic versus aerobic conditions (Fig. 2B).

**Figure 2.**
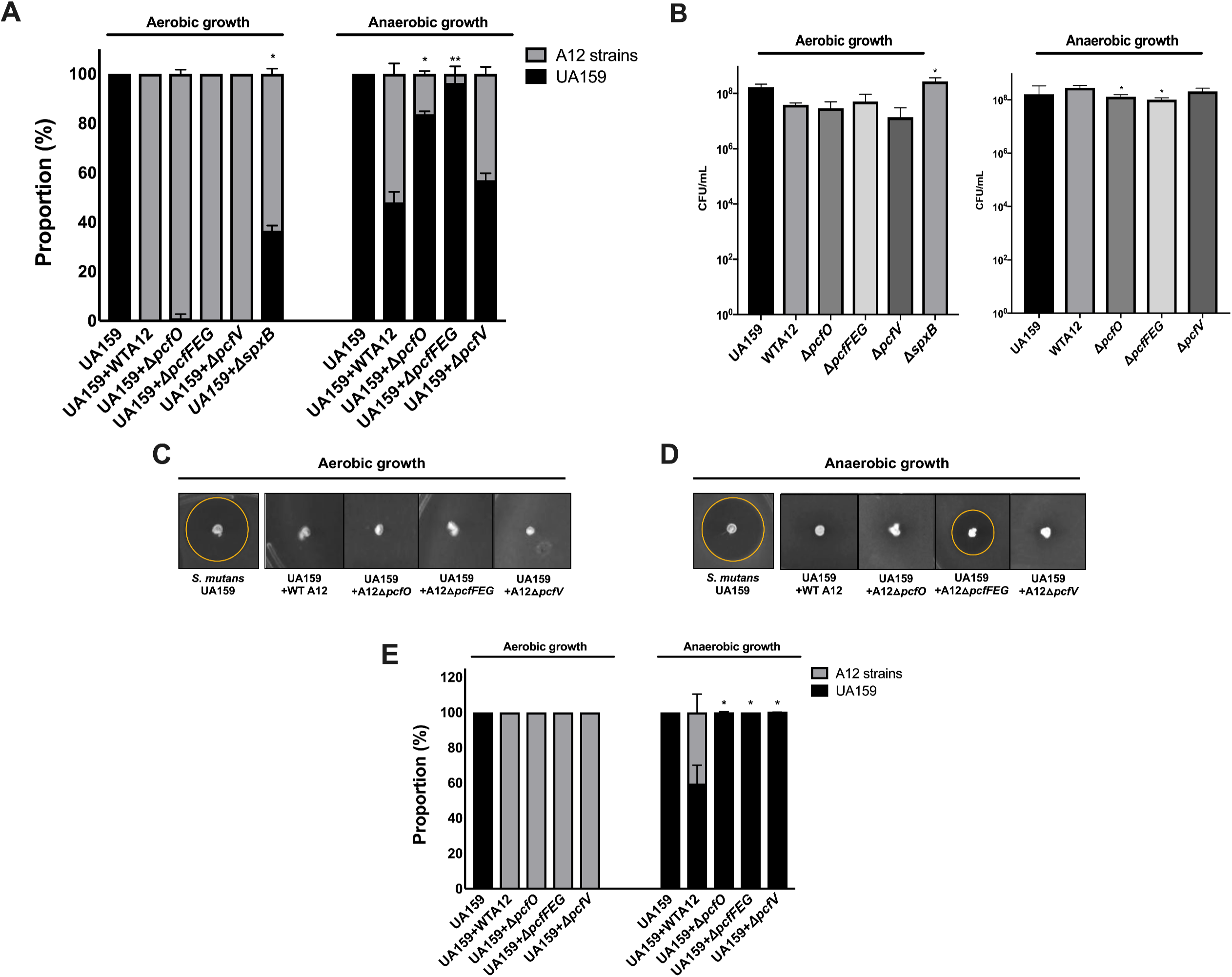
Impact of A12 <u>p</u>otential <u>c</u>ompetitive <u>f</u>actor. (A) Cultures of *S. mutans* UA159 (Sp^r^), A12, and A12 mutant strains were grown overnight and centrifuged to collect the cells. The cells were then washed with PBS and cultures were adjusted to OD_600_ = 0.5 using PBS. *S. mutans* was then mixed with A12 or A12 mutant strains at a ratio of 1:1. Mixed cultures were spotted on BHI plates with a sterile toothpick and incubated aerobically (5% CO_2_) or anaerobically at 37°C for 24 h. Pure cultures (B) were spotted on BHI plates with sterile toothpick as a control. (C,D) Mixed cultures of S. mutans UA159 (Sp^r^) and A12 or A12 mutant strains adjusted to OD_600_ = 0.5 were spotted on BHI plates and grown aerobically or anaerobically at 37°C. After 24 h, 3 ml of soft agar overlay (1% BHI agar) was mixed with 10^7^ cells of the indicator strain *S. sanguinis* SK150 and poured evenly onto the plate. All plates were incubated for an additional 24h. (E) *S. mutans* and A12 cells were enumerated by plating and the proportions of each (*S. mutans*, dark bars; A12, light bars) were calculated. Assays were performed at least three times, each performed with technical triplicates. Values are averages and error bars indicate standard deviations. Asterisks indicate statistically significant differences in proportions of *S. mutans* present when cocultured with A12 mutant strains compared with *S. mutans* cocultured with wild-type A12. In control assays, asterisks indicate statistical differences in CFUs of A12 mutant strains compared to CFUs of wild-type A12. Statistical analysis was performed using an unpaired Student’s t-test; * P < 0.05, ** P < 0.01.

A12 is able to interfere with the ability of *S. mutans* to produce bacteriocins using a protease encoded by the *sgc* gene that apparently degrades CSP, thereby blocking activation of bacteriocin genes by the ComDE two-component system (22). To evaluate whether the identified genes in A12 influence the ability of A12 to modulate the production of bacteriocins by *S. mutans*, a modified deferred antagonism assay was performed with the dual-species, plate-based cultivation system described above, where equivalent numbers of *S. mutans* and A12 or its derivatives were stabbed onto an agar plate. After 24 hours of incubation of the plates in aerobic or anaerobic conditions, 10^7^ cells of *S. sanguinis* SK150, an indicator strain that is sensitive to the mutacins produced by *S. mutans* UA159, were overlaid in soft BHI agar and plates were incubated for an additional 24 hours. As previously observed for aerobic conditions, A12 strains dominated the population, so growth inhibition of *S. sanguinis* SK150 by *S. mutans* mutacins did not occur, presumably because insufficient *S. mutans* survived to produce inhibitory quantities of mutacins (Fig. 2C). Interestingly, under anaerobic conditions, the A12 mutants lacking *pcfO* or *pcfV* behaved similarly to wild-type A12 (Fig. 2D), in that these mutant strains inhibited the ability of *S. mutans* to kill *S. sanguinis* SK150, despite the fact that the co-cultures were dominated by *S. mutans* (Fig. 2E). In contrast a zone of inhibition was observed when *S. mutans* was co-inoculated with A12Δ*pcfFEG*, albeit smaller than when *S. mutans* alone was inoculated (Fig. 2D). As was the case for the *pcfO* and *pcfV* mutants, *S. mutans* dominated the population when A12Δ*pcfFEG* was co-inoculated with *S. mutans.* Thus, deletion of *pcf* genes resulted in a markedly diminished capacity of A12 strains to kill *S. mutans* and/or to tolerate antagonistic factors of *S. mutans*.

### PcfO interferes with XIP signaling by *S. mutans*

Natural genetic competence is considered an important trait for *S. mutans*, not only because it is a primary driver for genetic diversification (48), but also because competence overlaps with regulatory circuits controlling multiple virulence-related traits, including bacteriocin production, biofilm formation and acid tolerance (30, 49, 50). While the CSP-ComDE circuit is essential for bacteriocin production and can affect induction of the competent state in *S. mutans*, the regulatory system directly controlling activation of competence is ComRS, with ComS serving as the precursor to the XIP (*com<u>X</u>*-inducing <u>p</u>eptide) signaling peptide. Culture supernates of A12 are apparently able to degrade XIP, which is not the case for *S. gordonii* (22) and some other commensals (51). Further, deletion of *sgc* in A12 did not diminish the ability of A12 supernatant fluids to block XIP-dependent activation of *comX*. The A12 genome contains a gene for a putative neutral endopeptidase that shares a substantial degree of homology to proteases of the neprilysin (M13) family, comprised of evolutionary conserved zinc-metallopeptidases showing functional diversities in numerous physiological processes (52). Members of the M13 protease family exist in a wide range of organisms, including mammals and bacteria, and are recognized as important regulators involved in the processing and degradation of a variety of different signaling peptides (52, 53). Having 71% identity (82% similarity) to PepO of *S. pyogenes*, which has been reported to degrade small hydrophobic peptides and disrupt quorum sensing (47), A12 PcfO was hypothesized to encode a PepO-like protease that could degrade XIP. Culture supernates were collected from overnight cultures of A12, A12Δ*pcfO*, *S. gordonii* DL1 or *S. mutans* UA159 grown in FMC (29). Various concentrations of sXIP were added to filter-sterilized culture supernates and incubated overnight, as detailed in the methods section. An *S. mutans* strain carrying a *lacZ* gene fusion to the *comX* promoter was then mixed with the supernates and incubated for 2 hours. Cells were collected and β-galactosidase assays were performed. As previously reported, A12 supernates eliminated the ability of sXIP to activate *comX* expression, whereas robust XIP signaling was observed with *S. gordonii* DL1 supernates, comparable to levels seen with the *S. mutans* control supernates (Fig. 3A). Interestingly, deletion of *pcfO* in A12 effectively eliminated the ability of A12 supernates to inhibit XIP signaling. There is crosstalk in the signaling pathways that control bacteriocin expression (ComDE) and genetic competence (ComRS) in *S. mutans* (54), so to assess whether A12 PcfO affected CSP signaling, the ability of supernates from A12 strains to degrade CSP was assessed using BHI medium and a derivative of *S. mutans* carrying a *cipB* promoter fused to *lacZ* to monitor CSP-dependent signaling through ComDE. The *pcfO* mutant strain retained the capacity to interfere with CSP-dependent activation of *P_cipB_-*lacZ at a level similar to supernates from wild-type A12, consistent with the findings that Sgc is primarily responsible for degradation of CSP (Fig. 3B).

**Figure 3.**
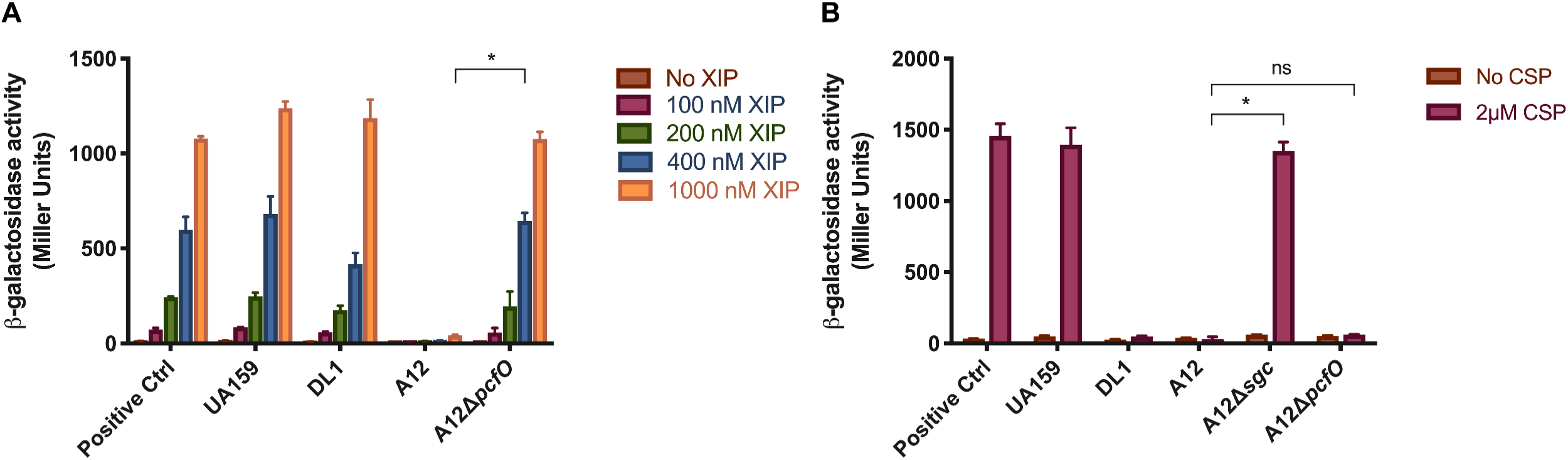
Effects of *S. gordonii* or A12 supernates on P*_comX_* or P*_cipB_* promoter activity. *S. mutans* containing a *lacZ* fused to the promoter of *comX* (A) or *cipB* (B) was grown to OD_600_ = 0.15 in FMC medium (A) or BHI medium (B). Supernates from overnight cultures of the indicated strains were obtained after centrifugation, the pH was adjusted to 7.0 with 6N NaOH and the solutions were filter sterilized. In the case of FMC conditions, supplemental glucose was added to increase the concentration of glucose by 25 mM. Synthetic peptides (XIP or CSP) were added to the supernates and incubated overnight (for XIP) or 2 hours (for CSP). The reporter strains were grown to OD = 0.15, cells were collected by centrifugation and then were resuspended in the indicated supernates for 2 h prior to measuring β-galactosidase activity. All assays were performed at least three times, with technical triplicates. The bars are averages and error bars represent standard deviations. Asterisks indicate statistically significant differences (A) between wild-type A12 (400 nM XIP) and A12Δ*pcfO* (400 nM XIP), and (B) between wild-type A12 and A12Δ*sgc* or A12Δ*pcfO*. Statistical analysis was performed using an unpaired Student’s *t*-test. * P < 0.05.

### PcfFEG contributes to immunity to the lantibiotic nisin

Bacteriocins appear to be highly influential ecological factors that enhance establishment and persistence of the producing bacteria. Bacteriocin-producing strains and many non-producing strains have immunity systems to provide protection against bacteriocins produced endogenously or by competing species (55). Because the identified PcfFEG ABC transporter shared homology with a lantibiotic immunity transporter using Blast searches, the susceptibility of A12 to the well-characterized and commercially available lantibiotic nisin was assessed. Overnight cultures of A12 were inoculated into fresh BHI medium in the absence or presence of various concentrations of nisin, and growth was monitored over the course of 24 h (Fig. 4A). Nisin at 2 µg/ml had no effect on the growth of A12. In the presence of 20 or 64 µg/ml nisin, the duration of the lag phase was substantially increased and the final optical density achieved was lower than without nisin, but the exponential growth rates were similar (66.5 ±2.75 min). To assess whether PcfFEG could provide protection against nisin, the growth characteristics of the Δ*pcfFEG* strain were compared with the parental strain in the presence or absence of lantibiotic. Additionally, because these types of ABC transporters can be regulated by two-component systems (TCS), the genes for the TCS (*pcfRK*; ATM98_05525 and ATM98_05520) encoded directly downstream of *pcfFEG* were deleted to create the Δ*pcfRK* strain of A12. Compared to A12, both Δ*pcfFEG* and Δ*pcfRK* displayed a significantly greater lag phase, even with 2 µg/ml nisin, than the wild type. Further, in the presence of nisin, the exponential phase growth rates of the mutants were slower and the final yields were decreased, compared to the parental strain. To confirm that *pcfFEG* expression could be influenced *pcfRK*, the *pcfFEG* were measured using qRT-PCR. As expected, in the absence of *pcfRK*, a significant decrease in the expression of *pcfFEG* was observed (Fig. 4B). Now, to directly examine the effects of nisin on the expression of the genes for the transporter and TCS, overnight cultures of A12 were sub-cultured in fresh BHI medium to OD_600_ = 0.5, then the cultures were treated with or without 2 µg/ml of nisin for 15 minutes (Fig. 4C). Treatment with nisin increased the expression of both the *pcfRK* and *pcfFEG* 4- to 6-fold, adding support to the idea that the function of these gene products is to respond to and confer protection against molecules that may be functionally and structurally similar to nisin. In fact, we have shown that as little as 6 nM nisin is sufficient to activate *pcfFEG* expression in a strain with intact *pcfRK* genes (data not shown).

**Figure 4.**
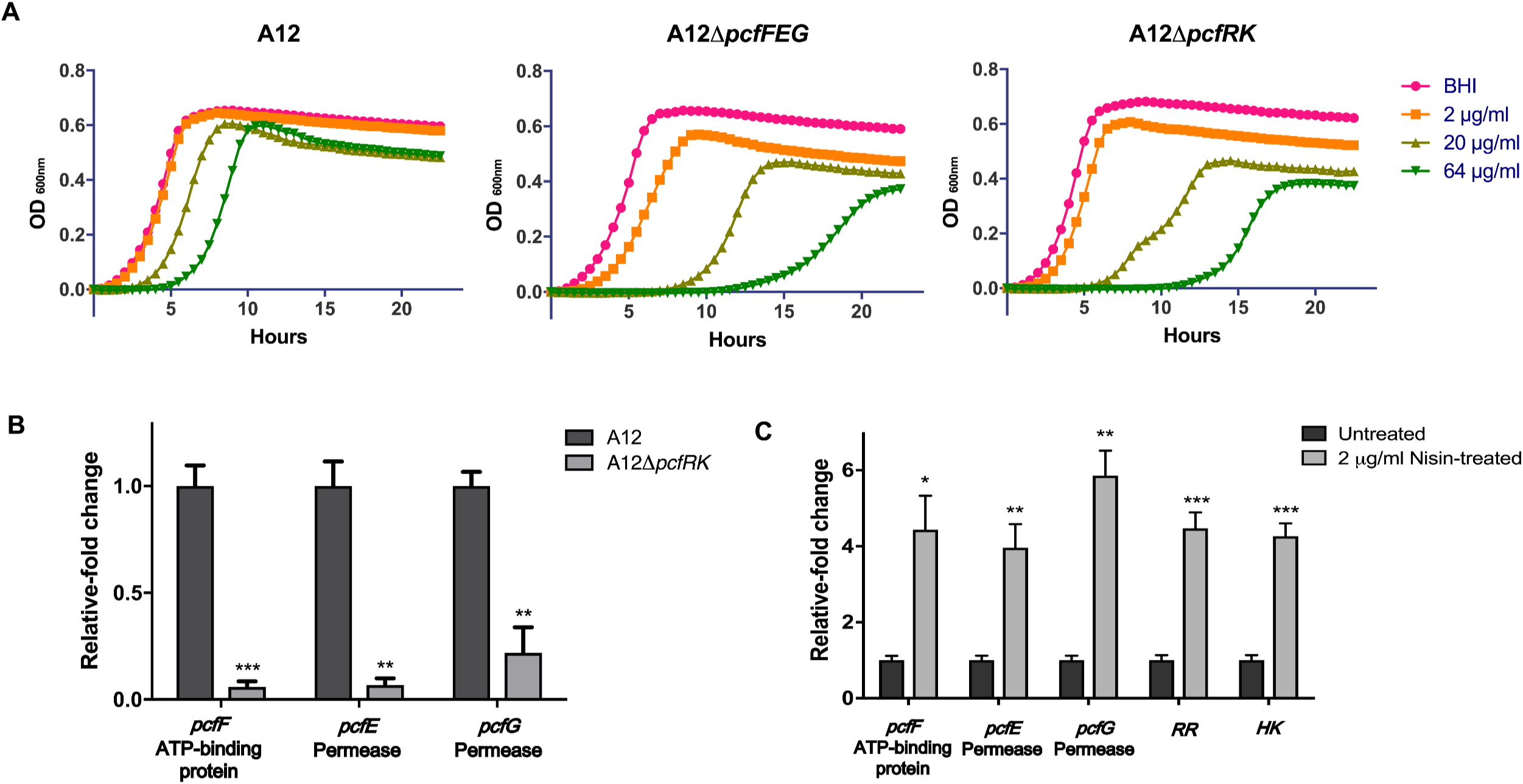
Deletion of *pcfFEG* or *pcfRK* increases susceptibility of A12 to the lantibiotic nisin. (A) Tolerance of A12 strains to different concentrations of nisin measured by monitoring growth OD_600_ over a 24-h period. Overnight cultures were subcultured into fresh BHI medium. Once the OD_600_ reached 0.5, cultures were diluted 1:100 into fresh BHI medium containing various concentrations of nisin and growth was monitored in a Bioscreen C. (B) Expression of *pcfFEG* mRNA was measured using qRT-PCR in the wild-type and *pcfRK* strain. Overnight cultures were inoculated into fresh BHI medium and once cultures reached OD_600_ 0.5, cells were collected for RNA extraction and subsequent qRT-PCR. (C) Expression of *pcfFEG* and *pcfRK* in A12 in response to treatment with 2 µg/ml nisin. RNA was extracted from mid-exponential phase cells of A12 treated with or without nisin, for subsequent qRT-PCR. All experiments were done in biological triplicates with technical triplicates. Error bars represent standard deviation and statistical analysis was performed using an unpaired Student’s t-test. * P < 0.05; ** P < 0.01, *** P < .001.

### Genetic elements involved in biogenesis of a putative antimicrobial molecule

As mentioned, bacteriocins play important roles in mediating competition among bacterial members of the same ecological niche. Vast diversity in composition, structure and mechanisms of action of bacteriocins have already been described, while genome sequencing and metabolomic approaches continue to identify new molecules (56–59). Colicin V is a small, secreted, pore-forming antimicrobial peptide produced by *E. coli* and its production involves several genes, including a *cvpA, cvaA* and *cvaB*; which encode proteins required for transport of colicin V across the cytoplasmic membrane with the help of TolC and an immunity protein, encoded by *cvi* (60). Additionally, the product of the *cvpA* gene is involved in the production of colicin V, although the precise function of its gene product is not established (61). Interestingly, the A12 genome contains a gene annotated as “colicin V production protein” (designated here as *pcfV*) that is encoded immediately downstream of an unannotated 96-aa protein (A12 peg.1179; ATM98_05820) and that is co-transcribed with *pcfV* (data not shown). To determine if there was a connection between the 96-aa ORF and *pcfV*, RNA was extracted from A12 and A12Δ*pcfV* and, after cDNA synthesis, qRT-PCR was performed. In the absence of *pcfV*, a 3-fold increase in the expression of the gene for the small hypothetical protein A12 peg. *1179* was observed (Fig. 5A), compared to A12. Next, the hypothetical protein was deleted and the mutant strain A12Δ*1179* was constructed to compare its behavior with that of the *pcfV* mutant. Both A12Δ*pcfV* and A12Δ*1179* showed decreased antagonism of *S. mutans*, when compared to the wild-type strain (Fig. 5B). Further, (Fig. 5A), deletion of *pcfV* resulted in a 3-fold increase in the expression of A12 peg. *1179.* However, deletion of A12 peg. *1179* did not affect the expression level of *pcfV* (p=0.049).

**Figure 5.**
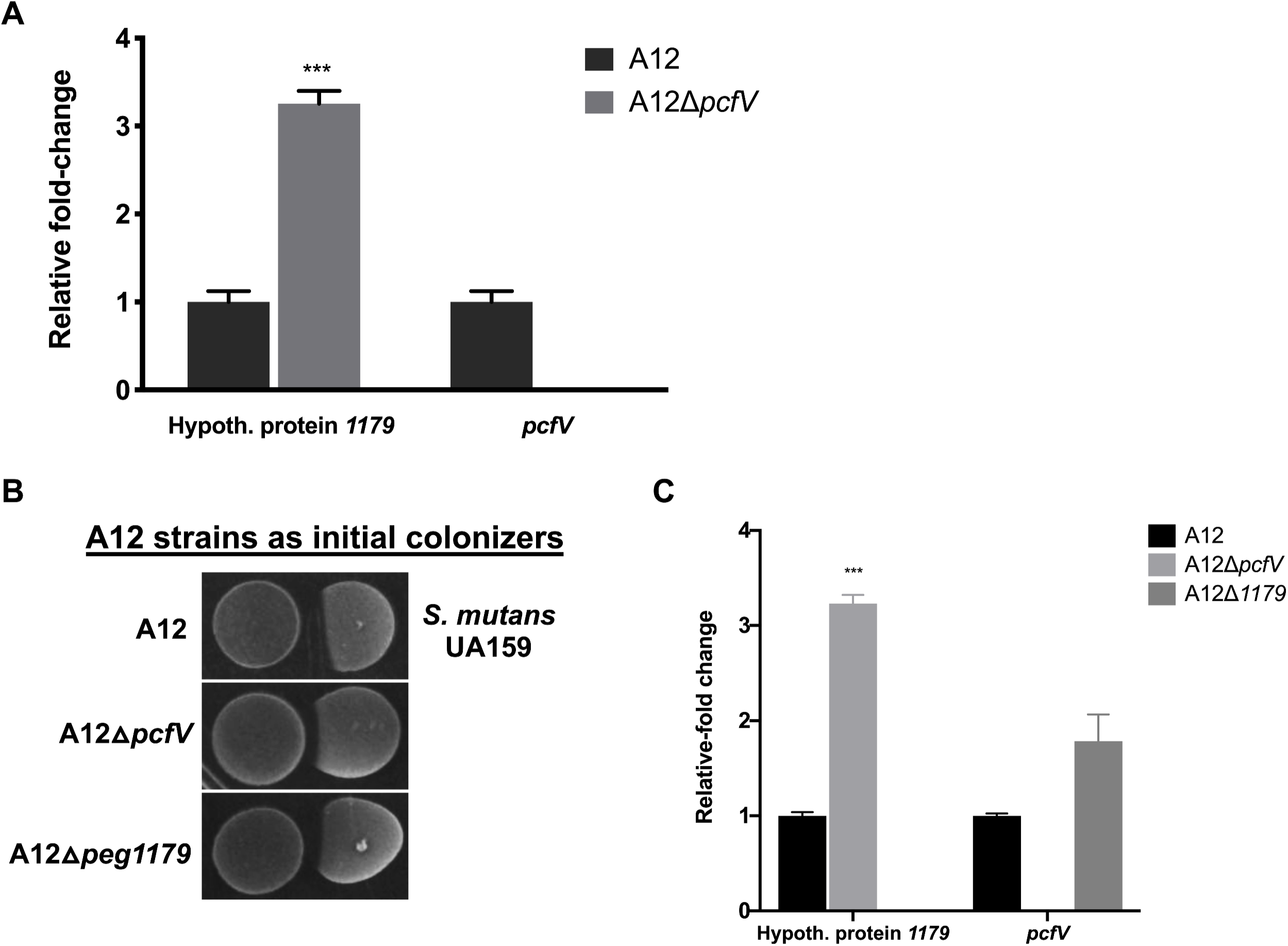
Expression of a hypothetical protein upstream of *pcfV* is up-regulated in the absence of *pcfV*. (A) Expression of *peg.1179* mRNA was compared using qRT-PCR in a *pcfV* deletion strain of A12 and the wild-type. Overnight cultures were inoculated into fresh BHI medium. When cultures reached an OD_600_ of 0.5, cells were collected for RNA extraction and subsequent qRT-PCR. (B) Plate-based growth inhibition of A12 strains versus *S. mutans*. All bacterial cultures were grown overnight in BHI medium, then adjusted to OD_600_ of 0.5 with fresh BHI. Aliquots (6 µl) from A12 or A12 mutant derivatives were spotted first, followed by spotting of *S. mutans* 24 h later. Plates were incubated for 48 h at 37°C 5% CO_2_ aerobic atmosphere. (C) Expression of *pcfV* of A12 was assessed using qRT-PCR in a *peg.1179*-deleted A12 strain and in the wild-type strain. All experiments were done in biological triplicates with technical triplicates, and error bars represent standard deviations. Statistical analysis was performed using an unpaired Student’s t-test; *** P < .001.

### Phylogenetic analysis of the *pcf* genes in other A12-like and closely related clinical isolates

Based on comprehensive phylogenomic analyses, the closest relative of the novel streptococcus isolate A12 is *S. australis* (22). Isolates that could not be appropriately classified as *S. australis* or *S. parasanguinis*, (another close relative of A12) due to significant differences in their genomes, are designated as “A12-like isolates”. To determine whether the identified *pcf* genes are present and conserved in strains that are closely related to A12, the genomes of four, low-passage *S. australis/*A12-like isolates were evaluated. In our analyses, the genome of *S. australis* ATCC700641 deposited in NCBI was also included, as were the genomes of a low-passage isolate *S. parasanguinis*, A1 (28), and *S. parasanguinis* ATCC903. A phylogenetic tree (Fig. 6A) shows the overall relationship of A12 to the other strains used for comparison, and the Roary matrix depicts the presence and absence of core and accessory genes of all isolates listed (Fig. 6B). The isolate *Streptococcus* G1 was more closely related to *S. australis*, whereas the *S. parasanguinis* reference strain and A1 were shown to be significantly different from A12, from strains designated as A12-like, and from *S. australis.* Next, A12 *pcf* sequences were used to do a series of BLASTP searches among the genomes of the strains of interest. ClustalW2 sequence alignment was done using the protein sequences obtained from strains to compare the percent identity against *Streptococcus* A12. All analyzed isolates encoded apparent PcfO and PcfV proteins, although some sequence variation was evident. Interestingly, the PcfV sequences in isolates closely related to A12 were highly conserved, with 100% identity to that of A12, whereas the *S. australis* ATCC700641 PcfV sequence showed 97.3% identity, and *S. parasanguinis* A1 and ATCC903 PcfV sequences were only 58.7% identical to A12 PcfV (Fig. 6C). The percent identity pattern for the small hypothetical protein A12 peg. 1179 was also similar, with all A12-like isolates being 100% identical to A12 peg. 1179 (Fig. S3). PcfO was more conserved, with less sequence variability, among the different streptococci included here, consistent with what has been reported for PepO from other *Streptococcus* spp. (62). On the other hand, substantial sequence variations were found in PcfFEG proteins from the different strains, except for A13 and *S. parasanguinis* ATCC903, which were 100% and ∼96% identical to A12 PcfFEG, respectively. PcfFEG sequences in other isolates were also identified by Blast of A12 PcfFEG sequences against their genomes. Identified sequences were subsequently aligned, but very low degrees of identity (5.5% to 20.2%) were observed. Given the overall similarity between ATP binding cassette transporters (63), a conclusion about whether PcfFEG homologues are present in strains other than A13 cannot be reached at this time. Furthermore, PcfRK sequence alignment showed a similar pattern, where considerable sequence variations were observed among the strains, except for A13 (100% identical to A12) and *S. parasanguinis* ATCC903 (99.1% and 97.4% identical to RR and HK sequences of A12, respectively).

**Figure 6.**
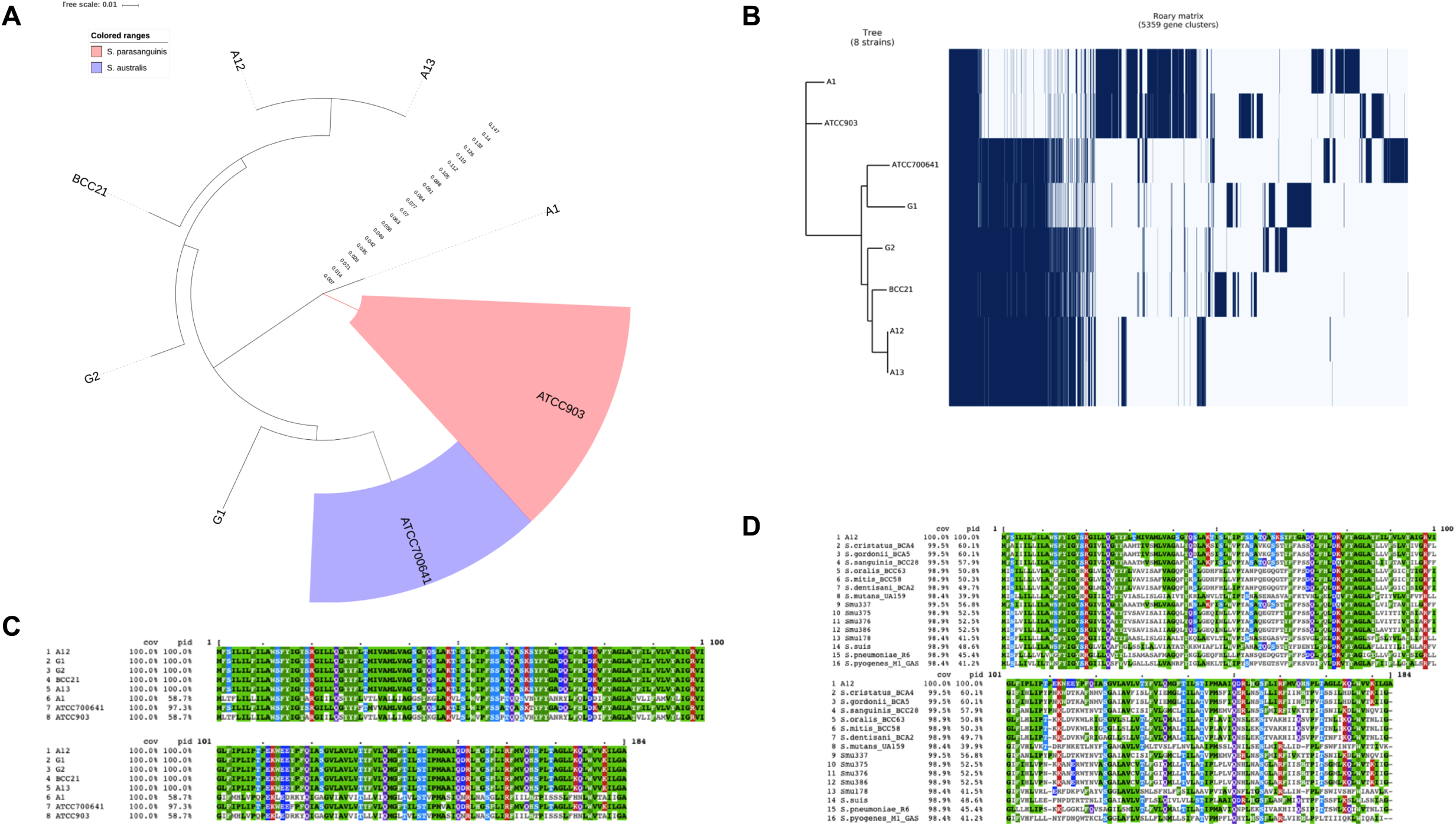
(A) Pangenome analysis of A12, A12-like isolates, *S. australis* ATCC 700641, and *S. parasanguinis* ATCC 903 (See text for details). (B) Presence and absence of core and accessory genes of all isolates listed. (C) CLUSTAL Omega alignment of *pcfV* in A12, A12-like isolates, *S. australis* ATCC 700641 and *S. parasanguinis* ATCC 903. Reference sequence designated as A12. % identity indicated as “pid”. (D) CLUSTAL Omega alignment of A12 *pcfV* amino acid sequences in other oral streptococci species.

Because PcfV was highly conserved in A12-like isolates, we explored whether *pcfV* genes were present in other oral streptococci. For this analysis, the genomes of 607 streptococci (26, 28) were analyzed. PcfV was also used in BLASP searches against three non-oral streptococci species (*S. suis, S. pneumoniae*, and *S. pyogenes*). Sequences similarity varied among and across species, ranging from 39% to 60% identity), suggesting that PcfV homologues/orthologues/paralogues were present in most streptococci.

## Discussion

Intermicrobial interactions in biofilms are central drivers of homeostasis and dysbiosis, and can have a powerful influence on maintenance of health or development of disease. An active role for commensal bacteria in providing protection against disease has been demonstrated in a variety of studies and environments, particularly for gut and skin infections (64–66). For example, a precisely defined consortium of commensal bacteria was reported to restore colonization resistance against vancomycin-resistant *Enterococcus* (VRE), a leading cause of nosocomial infection and a serious public health threat (67). This study showed that while the consortium of commensals worked cooperatively to resist VRE colonization, the previously uncharacterized bacterium *Blautia producta* was the key species within the consortium providing VRE-specific colonization resistance. Thus, the study of unique characteristics of certain key species in complex bacterial populations can shed considerable insights into the competitive and cooperative strategies that are employed by health-associated biofilms to persist and survive in particular ecological niches. Other examples of strain-specific inhibitory activities of commensals against pathogens are emerging (27, 68). Commensal oral streptococci, like A12, that can express high ADS activity under environmental conditions commonly occurring in oral biofilms, can directly inhibit growth of *S. mutans* by producing H_2_O_2_, and can disable bacteriocin production via interference with the CSP-ComDE pathway (22) may provide substantial protection to a human host against dental caries. Here, we demonstrate that multiple additional factors encoded by A12 are crucial to its ability to inhibit and to tolerate the most common human dental caries pathogen, *S. mutans*.

Bacteria in heterogenous biofilm environments are able to communicate through quorum sensing, which allows the coordination and regulation of genes crucial for survival (69). For the oral pathogen *S. mutans*, genetic competence is governed by the quorum-sensing peptides CSP and XIP. Here, we demonstrate that an endopeptidase of A12, encoded by *pcfO*, interferes with the *S. mutans* ComRS-XIP signaling pathway. In *S. mutans, comX* activation via ComRS enhances cell lysis (31), which results in increased eDNA release. The released eDNA has the potential to alter the structure of the biofilm matrix, which can impact the composition and pathogenic potential of the microbiota therein (70, 71). Additionally, autolysis of *S. mutans* is a primary mechanism for release of XIP, which can serve as a signal to other *S. mutans* to tune its competence system to assimilate exogenous DNA, providing nutritional benefits and contributing to the diversification of the species (31). It is also notable that the ComRS-XIP system in *S. mutans* is highly conserved among *S. mutans* isolates (26) and that ComR of *S. mutans* is specifically recognizes the XIP of *S. mutans*, but not XIP from other streptococci. Likewise, even though a *pepO* homologue is present in *S. gordonii* DL-1, there is no evidence that *S. gordonii* can interfere with XIP signaling in *S. mutans.* Given the conservation of PepO in *Streptococcus* species and its recently discovered role in virulence-related phenotypes in pathogens, it is possible commensal species like A12 have evolved to utilize the protease to enhance competitiveness in certain environments and/or against certain competing species. The ability of A12 to target a highly conserved and specific system of *S. mutans* imbues A12 with the ability to subvert key pathways of the pathogen and reduce the pathogenic potential of oral biofilm communities. Furthermore, the ability of A12 to degrade XIP using PcfO protease is particularly intriguing in light of the diverse functions of PepO being reported in *Streptococcus* species. For example, in addition to the ability of PepO to degrade small hydrophobic peptides (47), PepO has also been reported to affect a major virulence factor (SpeB) in *S. pyogenes* (72), mediate immune evasion in both *S. pneumoniae* and *S. mutans* (73–75), and even play a role in *S. mutans* UA140 mutacin production (62, 76). These recent findings on PepO highlight the complex roles of this peptidase in the fitness and physiology of various streptococci, including that of A12. It also raises the question of whether PepO proteins of A12 and other commensal oral streptococci may have the capacity to modulate the virulence of respiratory pathogens.

In addition to identifying the gene product that allows A12 to block XIP signaling in *S. mutans*, we establish here that there are multiple factors that are encoded by A12 that have a potent influence on the capacity of A12 to antagonize and to tolerate *S. mutans.* In aerobic conditions (Fig. 2A), the production of H_2_O_2_ masked the effects of deletion of *pcfO, pcfV, pcfFEG* on competition with *S. mutans*, but oxygen concentrations and redox conditions in dental plaque, particularly in mature dental plaque when conditions are favorable for caries development, are often sub-optimal for the generation of H_2_O_2_ by commensals. It is also clear from our data that, even when the ability of A12 to produce H_2_O_2_ is substantially impaired through the deletion of *spxB*, A12 is able to effectively compete with *S. mutans* (Fig. 2A); adding further support to the hypothesis that interspecies competition is a complex, multi-factorial process. In fact, under anaerobic conditions, when A12 is unable to produce H_2_O_2_, deletion of *pcfO, pcfFEG* or *pcfV* clearly compromises the capacity of A12 to compete with *S. mutans*. Furthermore, the loss of the *pcf* genes in A12 decreased the ability of A12 to persist and cope with antagonistic factors of *S. mutans.* For example, deleting *pcfFEG* in A12 impaired the ability of A12 to interfere with *S. mutans* mutacin production, largely due to the fact that the *pcfFEG* mutant constituted only about 4% of the viable counts recovered after 24 h of co-cultivation with *S. mutans*. Although the deletion of *pcfV* and *pcfO* did not impair the ability of A12 to interfere with bacteriocin production by *S. mutans*, significantly lower proportions of those A12 mutant strains were recovered after 48 h of co-cultivation. While the co-culture assays shown in Fig. 2A and 2E are performed with minor differences, the A12 mutant strains studied here consistently performed more poorly against *S. mutans* than wild-type A12.

Various commensal species have been shown to have inhibitory effects on *S. mutans*, including *S. gordonii, S. salivarius* and *S. dentisani* (77–79). Likewise, certain *S. salivarius* strains produce a bacteriocin-like inhibitory substances (80, 81) that have made them potential oral probiotic candidates. While commensals may directly antagonize pathogens via the secretion of antimicrobial peptides, our findings suggest that commensals may also have genetic elements that allow them to mount a defense against other members of complex oral microbial communities. Deletion of *pcfFEG* in A12 resulted in a diminished ability to compete with *S. mutans* and our early exploration into the mechanism of this genetic locus revealed a role in immunity to nisin, a well-characterized lantibiotic. Lantibiotics are a class of bacteriocins produced by Gram-positive bacteria that undergo post-translational modification and contain either lanthionines or methyl-lanthionines ring structures with atypical amino acids (82). The producing organisms usually encode specific immunity proteins that provide protection against the deleterious effect of their own lantibiotics. Two immunity systems have been described (83), one being a dedicated ABC transporter thought to prevent accumulation of lantibiotics in the cell membrane. The second type involves lipid-anchored immunity proteins that are believed to sequester lantibiotics at the cell surface before they can cause damage (83). Various strains of *S. mutans* produce bacteriocins (called mutacins) that belong to the class of lantibiotics, including mutacins I, II, III, K8, B-Ny266, Smb, and 1140 (76). However, not all strains of *S. mutans* produce lantiobiotic-type mutacins. Notably, *S. mutans* UA159 encodes only non-lantibiotic mutacins (84). Still, we observed a reduced persistence of A12 lacking PcfFEG when co-cultured with *S. mutans* UA159. Thus, our hypothesis is that PcfFEG of A12 can function not only as a system that can be induced by, and confer protection against, lantibiotics, but that PcfFEG may confer resistance of A12 to multiple antimicrobial peptides. There is precedent for broad specificity in immunity proteins, with one example being the SmbFT system of *S. mutans* GS5, which provides protection against a number of different, but structurally similar lantibiotics (85). Further characterization of the PcfFEG system is ongoing to understand what types of molecules can induce the system through the cognate TCS identified here (PcfRK) and how resistance profiles to different antimicrobial peptides are affected by the PcfFEG proteins; with the overarching goal of understanding how PcfFEG can enhance the fitness of A12 in complex microbial communities.

Another novel finding reported here is that A12 harbors a gene that is homologous or orthologous to a putative colicin V biosynthetic protein; *pcfV*, which is similar to *cvpA* in *E. coli*. Although the role of *cvpA* in Colicin V biogenesis of *E. coli* is not particularly well understood, the production of mature Colicin V, which is ribosomally-synthesized from a small precursor protein, is thought to require CvpA (61). Furthermore, while colicins were first studied in *E. coli*, they have also been found to have similarities to bacteriocins of Gram-positive bacteria (86). More recently, high-throughput screening for antimicrobial activity against *Staphylococcus aureus* on coagulase-negative commensal staphylococci isolated from the skin of healthy individuals disclosed previously unknown antimicrobial peptides (AMPs) encoded within the genomes of the isolates exhibiting active antimicrobial activity; among the genes identified was one annotated as “colicin V production protein” (87). Interestingly, A12 *pcfV* is similar in size and has 30 to 40% sequence identity to those identified in commensal *Staphylococcus* species. Here we found that the *pcfV* mutant of A12 has diminished capacity to antagonize *S. mutans*. Notably, we also show that the expression of a small hypothetical protein encoded upstream of *pcfV* can be altered by deletion of *pcfV* and that deletion of only this peptide confers a phenotype that is similar to what is observed in a strain lacking PcfV. Our working hypothesis is that this peptide may serve as a precursor that is acted on by PcfV and possibly other gene products to produce a mature antimicrobial peptide that may inhibit growth or metabolism of *S. mutans*, and possibly other organisms in the oral microbiome. It is noteworthy that A12 encodes a gene product annotated as colicin E2 immunity protein, albeit at a site distant from *pcfV*. Additional studies to identify the product of PcfV and determine whether the predicted colicin E2 immunity protein is related to PcfV function or involved in protection of A12 against a colicin-like antimicrobial factor(s) are underway.

We have only begun to uncover the diverse mechanisms utilized by beneficial, health-associated organisms to persist against pathogens. Work is in progress to build a more comprehensive understanding of the interaction between A12 and *S. mutans* or other pathogens, and how the identified *pcf* genes may influence the expression of other genes related to inter-species competition by A12 or the virulence of *S. mutans.* As reported here, other A12-like isolates have similar phenotypic properties to A12 (51) and many of these A12-like isolates appear to encode the competitive factors studied here in A12. In fact, *pcfV* shows an exceptionally high degree of conservation in A12-like organisms and close relatives of A12, as well as being broadly distributed in oral streptococci. Another important point is that PcfFEG was only found in A12 and A13, an isolate in the group of “A12-likes”. The presence and absence plot (Fig. 6B) suggests that A12 and A13 are most similar in their gene content. If *pcfFEG* can indeed confer protection against a broad range of bacteriocins, perhaps A12 and other A12-like streptococci acquired and retained these genes to be able to persist in the hostile environment of the oral cavity. Overall, then, the mechanisms we have identified begin to provide substantive new insights into the complexities of interbacterial competition in the oral microbiome, and contribute to the foundation of knowledge needed to understand how beneficial commensal species promote oral health. Continued genome-scale analysis, coupled with functional genomics and metatranscriptomics using the expanded database of sequence of commensal organisms (28), will not only help to understand the forces that shape and maintain the oral microbiome, but the information can be used for the rational design of novel therapies that employ pre-, pro- or synbiotic approaches to control oral and other infectious diseases.

## ACKNOWLEDGMENTS

This work was supported by the National Institute of Dental and Craniofacial Research of the National Institutes of Health under Award Numbers R01 DE025832, T90 DE021990, and F30 DE028184.

